# Base editing using CRISPR/Cas9 in *Drosophila*

**DOI:** 10.1101/2021.03.24.436868

**Authors:** Elizabeth Marr, Christopher J. Potter

**Affiliations:** Solomon H. Snyder Department of Neuroscience, Johns Hopkins University School of Medicine, 855 N. Wolfe St, Baltimore, MD, 21205, USA; Mercy Medical Center, 345 St. Paul Pl, Baltimore, MD 21202, USA

**Keywords:** base editing, BE2, mutator, CRISPR/Cas9, cytidine deaminase, mutator, Drosophila melanogaster, insect, guide RNAs

## Abstract

Cas9 and a guide RNA function to target specific genomic loci for generation of a double stranded break. Catalytic dead versions of Cas9 (dCas9) no longer cause double stranded breaks and instead can serve as molecular scaffolds to target additional enzymatic proteins to specific genomic loci. To generate mutations in selected genomic residues, dCas9 can be used for genomic base editing by fusing a cytidine deaminase to induce C>T (or G>A) mutations at targeted sites. Here, we test base editing in *Drosophila* by expressing a transgenic *Drosophila* base editor (DBE2, based on the mammalian BE2) which consists of a fusion protein of cytidine deaminase, dCas9, and uracil glycosylase inhibitor. We utilized transgenic lines expressing gRNAs along with pan-tissue expression of the *Drosophila* Base Editor (*Actin5C-DBE2*) and found high rates of base editing at multiple targeted loci in the 20 bp target sequence. Highest rates of conversion of C>T were found in positions 3-9 of the gRNA targeted site, with conversion reaching nearly 100% of targeted DNA is somatic tissues. The simultaneous use of two gRNA targeting a genomic region spaced ∼50 bps apart led to mutations between the two gRNA targets, implicating a method to broaden the available sites accessible to targeting. These results indicate base editing is efficient in *Drosophila*, and could be used to induce point mutations at select loci.

## Introduction

The development of clustered regularly interspaced short palindromic repeat (CRISPR)/ Cas9 as a method to target specific genomic loci has revolutionized genetic engineering (Anzalone et al., 2020). Cas9 is a homing endonuclease that uses a small RNA molecule to target and then cut DNA. Endogenous Cas9 binds to a complexed pair of small RNAs (crRNA and tracrRNA) that combined form a single RNA commonly referred to as a single guide RNA or guide RNA (gRNA or gRNA) (Brouns et al., 2008; Cong et al., 2013; Jinek et al., 2013; Mali et al., 2013). The typical two-component system for CRISPR/Cas9 thus minimally requires the Cas9 endonuclease to be complexed with a gRNA. The Cas9/gRNA complex can be reconstituted *in vitro* or expressed from transgenic constructs and formed *in vivo*. Guide RNAs typically include 19-20 base pair homology to the targeted sequence whose only restriction is that it must lie directly next to a protospacer-adjacent motif (PAM) DNA sequence (Gasiunas et al., 2012; Cong et al., 2013; Jinek et al., 2013; Mali et al., 2013). The PAM for *S. pyogenes* Cas9 is *NGG* (Jinek et al., 2013).

The Cas9/gRNA system has been used effectively in *Drosophila* for the generation of targeted double strand breaks in DNA (Bassett et al., 2013; Gratz et al., 2013; Kondo and Ueda, 2013; Ren et al., 2013; Yu et al., 2013). Two repair pathways are induced by double-strand breaks: non-homology end joining (NHEJ) or homology directed repair (HDR). If the double-strand break is repaired by non-homologous end joining, this often results in insertions or deletions (indels) at the targeted cut site, which can lead to the generation of null mutations in the targeted gene. Homology directed repair attempts to correct the double strand break by using homologous DNA surrounding the cut to copy in DNA that might have been lost at the break point. The homologous template is usually the sister chromosome, but an exogenous homologous template can also be provided containing a genetic cargo flanked by DNA arms homologous to those generated by the double strand break. This can be used to generate knock-ins at the genetic locus.

The adoption of CRISPR/Cas9 into the *Drosophila* toolbox has made it easy to generate null mutations for a gene of interest (Bassett et al., 2013; Gratz et al., 2013; Ren et al., 2013; Gratz et al., 2014). However, as demonstrated by genetic screens which utilized chemical mutagenesis (Nüsslein-Volhard and Wieschaus, 1980), point mutations in a gene can lead to a variety of alterations of gene function that reveal insights in how a gene product functions in a cell. They can, for example, reveal important residues in a functional domain, or identify interaction domains (Kaufman, 2017). A reliance on CRISPR/Cas9 as the predominant method for generating mainly null mutations could lead to a lack of point mutations in a gene.

Mutated versions of Cas9 have been developed that deactivate its endonuclease function (dCas9), yet still retain its ability to target specific genetic loci as directed by a gRNA (Bikard et al., 2013; Gilbert et al., 2013; Qi et al., 2013). The dCas9 thus serve as a homing scaffold to direct the targeting of other proteins to specific DNA sites, including proteins that can modify DNA. This approach has been used to convert dCas9 into a base editor to generate point mutations at a target site (Komor et al., 2016). In this approach, dCas9 is tethered to cytidine deaminase (CD), which directs the deamination of cytosine (C) residues to uracil (U) (Komor et al., 2016). Most cytidine deaminases use RNA as a substrate, but some cytidine deaminases can also utilize single-stranded DNA as a substrate (Komor et al., 2016). The Cas9/gRNA complex can lead to single-stranded DNA at the templated site, which can serve as a template for a tethered cytidine deaminase. Uracil has the base pairing properties of thymine and pairs with adenine. During repair or DNA synthesis, the uracil can lead to incorporation of an adenine at the template strand. This effectively converts cytosines in DNA to thymine. A number of base editors have been developed based on this strategy. Base editor 2 (BE2) tethers the rat cytidine deaminase APOBEC1 to the N-terminus of dCas9 and uracil DNA glycosylase inhibitor (UGI) to its C-terminus (**Figure 1A**). UGI acts to inhibit the natural DNA repair response to remove Uracil from DNA via uracil DNA glycosylase, and was found to increase base editing efficiency three fold in cultured cells (Komor et al., 2016). In cultured cells, BE2 primarily converted cytosine residues in a base editing window corresponding to positions 4-8 in the gRNA targeted region.

**Fig. 1.**
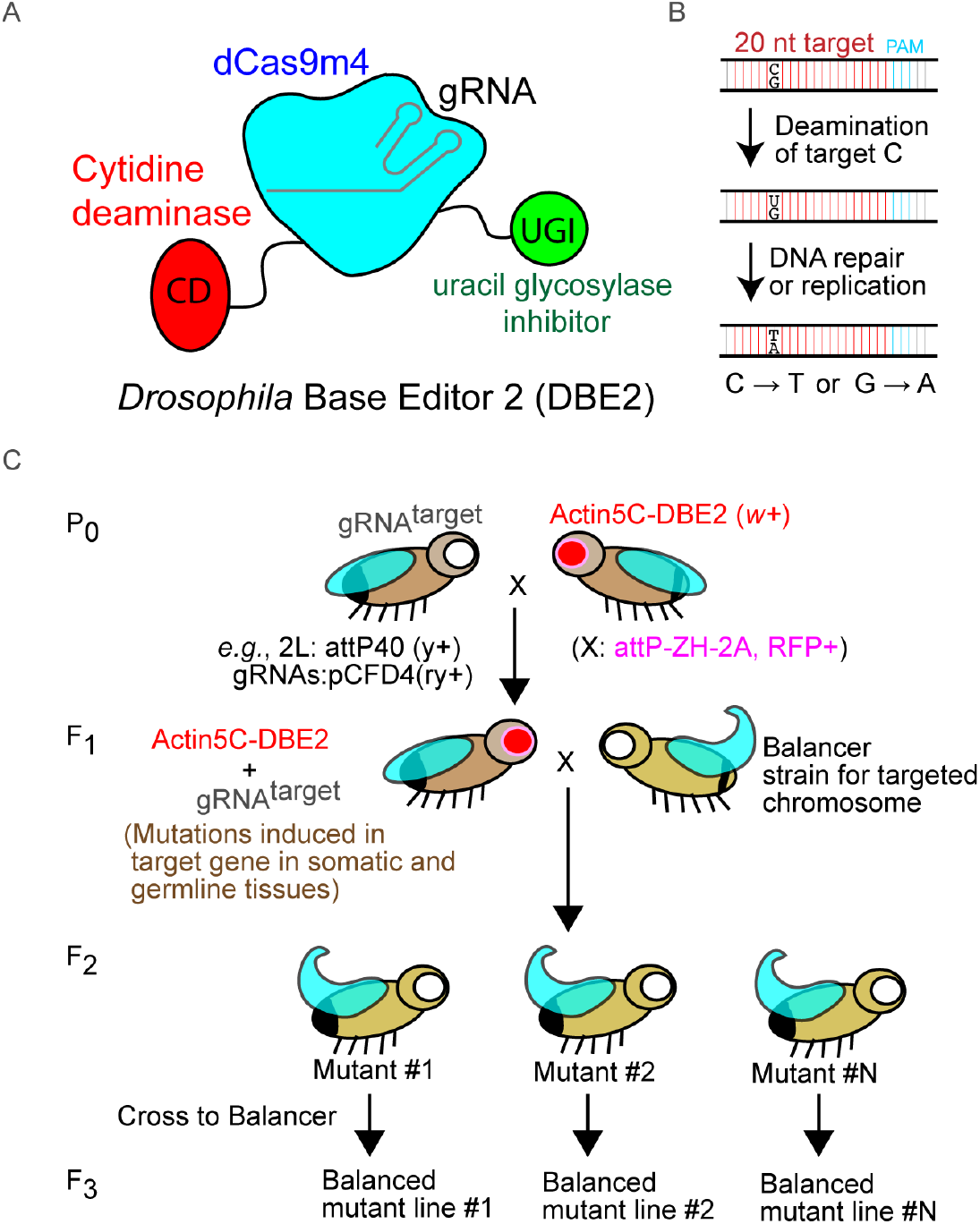
Base editing in Drosophila. A) Schematic of Drosophila Base Editor 2. A catalytically inactive dead Cas9 (dCas9m4) is fused via flexible linkers to cytidine deaminase and uracil DNA glycosylase inhibitor (UGI) domains. B) Summary of mechanisms inducing point mutations. A guide RNA (gRNA) directs the base editor to a 20 nucleotide target. A cytidine residue may be deaminated, resulting in conversion of the cytidine (C) to a uridine (U). During DNA repair or replication, the uridine has base-paring properties of thymine (T). This results in a C-to-T change on the top strand, or a Guanosine-to-Adenosine (G-to-A) change if the gRNA targeted the bottom strand. C) Summary of fly crosses used to induce base editing. In this example, males containing one or more gRNAs (inserted on the 2nd chromosome using the pCFD4 construct (Port et al., 2014)) are crossed to the base editor (inserted on the X chromosome) that uses the Actin5C promoter to express in all cells. Base editing occurs in male F1 progeny. Editing can be examined directly in F1 males, or mutant lines obtained by crosses to balancers to isolate the mutagenized chromosome.

Here we tested in *Drosophila* a *Drosophila* base editor (DBE2) similar in design to BE2. Expression of DBE2 in all tissues in combination with different transgenic gRNAs revealed robust and efficient conversion of cytosine residues to thymine across a base editing window of approximately 11 bases (position 1-11 of the gRNA target sequence). Conversions occured most frequently at positions corresponding to residues 3-9 in the gRNA target. Interestingly, the simultaneous use of two gRNAs targeting the same genomic locus but spaced approximately 50 bp apart broadened base editing to the region between the gRNA targets. The data presented here suggests base editing in *Drosophila* might be an effective method to induce point mutations at target genetic loci.

### Results and Discussion

The *Drosophila* base editor (DBE2) cassette was derived from Base Editor Version 2 (BE2) that consists of a fusion protein of Cytidine Deaminase, dCas9, and uracil DNA glycosylate (Komor et al., 2016). BE2 uses a dCas9 that contains the mutations D10A and J840A, referred to here as dCas9m2. In experiments using dCas9 as the scaffold for tethering transcriptional activator domains in *Drosophila*, the function of dCas9m2 was compared to dCas9m4, a variant of dCas9 that contains the same mutations as dCas9m2 with the additional mutations H839A and N863A. dCas9m4 was found to function more effectively as a tether for targeting loci in *Drosophila* than the dCas9m2 (Lin et al., 2015), although the reasons for this were unknown. Nonetheless, these results suggest that dCas9m4 may function as a better gRNA-targeted scaffold in *Drosophila* than dCas9m2, and as such, the dCas9m4 variant was used in place of the dCas9m2 variant found in BE2. This generated the *Drosophila* base editor used here (**Figure 1A**). DBE2 was cloned into an PhiC31 attB containing construct marked by mini-white for the generation of transgenic flies.

The Liu group also developed Base Editor Version 3 (BE3) in which the catalytic H839A mutation in BE2 was reverted, allowing the mutated Cas9 protein to function as a nickase and cut the non-edited guanine strand opposite the uracil. This increased the efficiency of base editing in cell culture by roughly 2 to 5-fold, but also led to insertion and deletions (indels) at the targeted site. So while BE3 was more efficient at inducing base editing compared to BE2, this came at the expense of also generating indels. When deciding to adapt BE2 or BE3 for base editing in *Drosophila*, we focused on the BE2 reagent in order to generate base editing without indels. We further reasoned that potential *in vivo* decreases in efficiency with BE2 could be easily compensated by increasing the number of *Drosophila* progeny screened. We also reasoned that potentially decreasing efficiency of BE2 might prove useful in future experiments attempting to isolate mutated flies containing a single targeted change.

To test the ability of the *Drosophila* base editor DBE2 to induce changes in *Drosophila*, the *Actin5C* enhancer was used to direct its expression in all tissues. Using PhiC31 integrase, the *pattB-Actin5C-DBE2* construct was inserted onto the X chromosome at site attP-ZH-2A (Bischof et al., 2007) which is marked by *3xP-RFP*. A stable homozygous stock of *Actin5C-DBE2* was established.

To target DBE2 to a variety of genomic loci in the *Drosophila* genome, the Harvard Transgenic RNAi Project collection of transgenic *U6:gRNAs* (Zirin et al., 2020) were used with *Actin5C-DBE2* (**Figure 1C**). This collection has been developed by the Perrimon lab to use transactivator variants of dCas9 to drive expression of endogenous genes (Chavez et al., 2015; Lin et al., 2015). The gRNAs in this collection targeting putative enhancer sites were cloned into the pCFD3 or pCFD4 vectors (Port et al., 2014), and integrated at the *attP40* location on the second chromosome. Thirty transgenic U6:gRNAs from this collection were used to test the ability of *Actin5C-DBE2* to induce mutations. The gRNAs selected for examination were chosen as mutation of the target genes were suspected not to cause lethal effects (which would interfere with the analyses). An example cross is shown in **Figure 1C**. Males containing the *U6:gRNA* transgene were crossed to virgins containing *Actin5C-DBE2* on the X chromosome. All male F1 progeny contain both components, and mutations will be induced in somatic and germline tissues. The advantage of using the *Actin5C* enhancer to direct expression is that F1 males can be directly examined for changes to somatic DNA. Mutant lines can be established by crossing these F1 males to a suitable balancer chromosome stock to isolate F2 males with the targeted chromosome over a balancer. A mutant line can be established in the F3 generation (**Figure 1C**).

In the experiments presented here, only F1 males were tested for changes in DNA. Future experiments will be required to determine the extent of germline transmission of the induced mutation using the *Actin5C* enhancer, or by expressing *DBE2* in the germline using appropriate enhancers such as *vasa* or *nos (Gratz et al., 2014; Port et al., 2014)*.

To determine if DBE2 can induce mutations in the genome, the targeted genomic region was PCR amplified in pools of ∼10 F1 adult males, and the target region sequenced by Sanger sequencing. To control for polymorphisms that might already be present at the targeted site, the genomic region of F1 males which used a gRNA targeting a different genomic locus (yet of the same genetic background) were also PCR amplified and sequenced by Sanger sequencing. Genomic loci that were found to contain multiple polymorphism at the targeted site, or in the gRNA target, were excluded from further analyses.

Of the 30 targeted locations, 15 demonstrated changes to their DNA. The gRNAs not resulting in changes are listed in *Materials and Methods*. All changes were C>T mutations as seen in the Sanger sequence chromatograms (**Figure 2A**) as expected by the function of cytidine deaminase. The cytosine residues showing changes in the chromatograms are shown in blue in **Figure 2A**. A summary of all genomic changes are summarized in **Figure 2B** in which cytosine residues that showed any changes in the chromatograms are highlighted in red. Mutations were not found outside this 20 base pair region. The majority of mutations occurred at positions 3-9 in the 20 base pair targeted region, although mutations were also found to occur at locations 17 and 19 (**Figure 2A, Figure 2B**), which was not reported in mammalian cell culture studies (Komor et al., 2016). These results are summarized in **Figure 2C**, which displays the frequency that any change was found at a cytosine residue at a certain location in the target. For example, the 15 gRNA targeted regions contained 5 examples in which a cytosine was located at position 8 in the targeted genomic loci, and 4/5 of these targeted sites (80%) demonstrated changes to that cytosine in the Sanger sequencing chromatograms. This gives an estimate of how often a particular cytosine residue at a certain location in the 20 nucleotide target site would be targeted by the DBE2. To estimate how efficiently this targeting might be occurring at each targeted site, the chromatograms were used with the EditR program (Kluesner et al., 2018) which calculates the percentage a residue is represented at each location. The chromatographs for all sequencing reactions were used to estimate the percentage of cytosine residues changed to a thymine at each location (**Figure 2D**). The cytosine residues at locations 3-9 were the most efficiently targeted, with some locations being targeted close to 100%, indicating that the somatic DNA in all the adult tissues had been mutated. For example, this is evident in the chromatographs for CG4998 (the cytosine at position 7 has been nearly converted to thymine), and for Cyp1 (the cytosine at position 5 is nearly all thymine). Future studies can extend these studies with additional gRNA from the TRiP collection to determine why certain locations are favored for mutation versus others. These data demonstrate that base editing in *Drosophila* tissues can occur at high efficiencies.

**Fig. 2.**
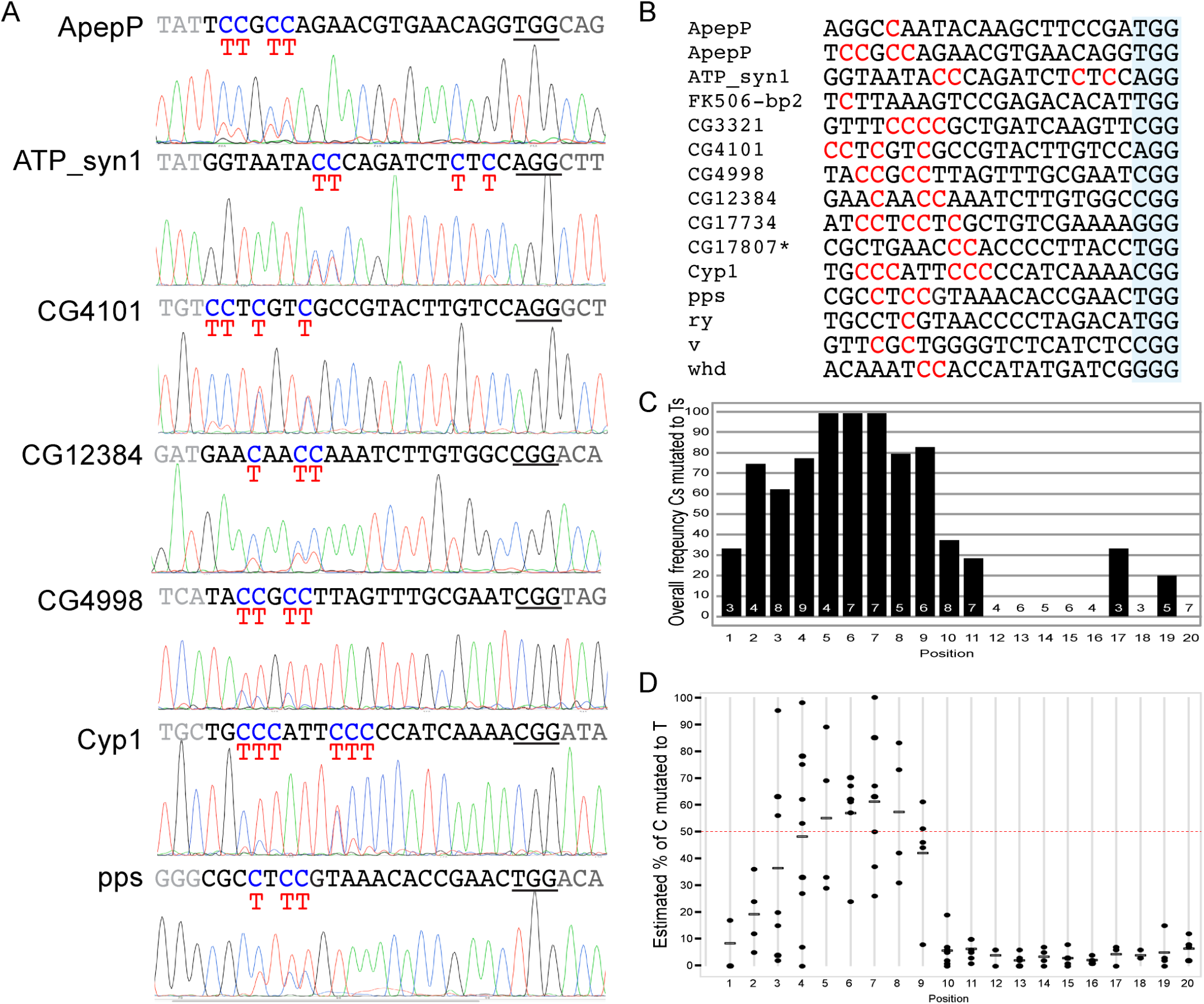
Example targets and efficiencies of base editor. **A)** The targeted genomic site of F1 male flies containing the base editor and a gRNA were PCR amplified and sequenced. Targeted cytosine (C) residues are shown in blue on top of the chromatographs. The PAM sequence is underlined. **B)** Fifteen different genomic loci were targeted by the base editor. Cytosine residues mutated by the base editor are shown in red. Cytosine residues not highlighted were not mutated. The PAM sequence is highlighted in blue. **C)** Examining the location of base editing within the gRNA target. The bar graphs shows the frequency that a cytosine residue at a particular location in the 20 base pair targeted site was mutated. The numbers in the bars indicate the number of times a cytosine was present in that position across all 15 targeted genes. For example, C>T mutations in positions 6-8 were found in all PCR-amplified targets examined from F1 flies. **D)** Estimating the efficiency of base edit targeting. The EditR program (Kluesner et al., 2018) was used with chromatographs to estimate the percentage of cytosine converted to thymine at each gRNA targeted site. Points above the red-dashed line at 50% suggest that both alleles were targeted. The line indicates the average.

The *Actin5C-DBE2* efficiently targeted cytosines in the 20 base pair targeted region. However, surprising results were found when a genomic region was targeted by two guide RNAs within ∼50 bps of one another. This was initially identified in experiments that used two gRNAs to target the GAL4 gene in transgenic *Drosophila* lines. The GAL4 gRNAs were located 52 base pairs apart, and had previously been validated to efficiently target GAL4 regions for knock-in using the HACK method and their effectiveness in inducing double stranded breaks and indels at these targeted GAL4 loci (Lin and Potter, 2016). The expectation for using these two guide RNAs would be the efficient induction of C>T changes in the two targeted locations. However, this was not found. Instead, cytosine or guanine residues (corresponding to changes of the cytosine on the bottom strand) between the gRNA targets were efficiently targeted (**Figure 3A**). The GAL4 targeted region, and changes to the DNA, are highlighted in red in **Figure 3A**. No mutations were found in the 20 base pair targeted sites, but were instead found between these targeted site (**Figure 3B**). The most frequently targeted site was a cytosine located 22 base pairs upstream of GAL4-gRNA1 on the bottom strand (as seen by a G>A mutation in the top sequence strand). This suggests that the DNA between these two gRNAs were now within the targeting window of the tethered cytidine deaminase. In addition, since cytidine deaminase requires single stranded DNA as a template, it suggest that the DNA stretch between the gRNAs targeted sequence might also be unpaired or single stranded. The efficiency of the targeting was estimated using the Sanger sequence chromatographs with the EditR program (**Figure 3B**). This confirmed that the bottom strand was being efficiently targeted (in some cases above 90% of all guanine residues at this site were converted to an adenine residue). Mutations also occurred at many locations in the region between the gRNA targeted loci, in particular a cytosine rich region. The ability of a base editor to expand its range when two gRNA were used to target a location has not to our knowledge been previously reported. To determine if this was specific to GAL4 sequences, or might be generalizable to other genomic locations, the available transgenic *U6:gRNA* collection from Bloomington was screened for gRNAs that targeted the top strand about 50 base pairs apart. Three gRNA pairs were found to match these criteria, and were tested with the *Actin5C-DBE2*. In all three cases, mutations were found in the region between gRNAs, although the efficiencies were lower than that found with GAL4 targeting (**Figure 3C**). The summary of the changes using these gRNA pairs with the EditR-estimated percentages, are shown in red in **Figure 3C**. Some of the gRNA targeted sites also demonstrated changes, in contrast to the GAL4 gRNA pairs. Overall, these data suggest that the use of two gRNAs can expand the targeted region of DNA which might not otherwise be compatible for Cas9 directed mutation. In addition, since the region undergoing base editing is outside the gRNA targeted region, repeated base editing might occur to completion in this area.

**Fig. 3.**
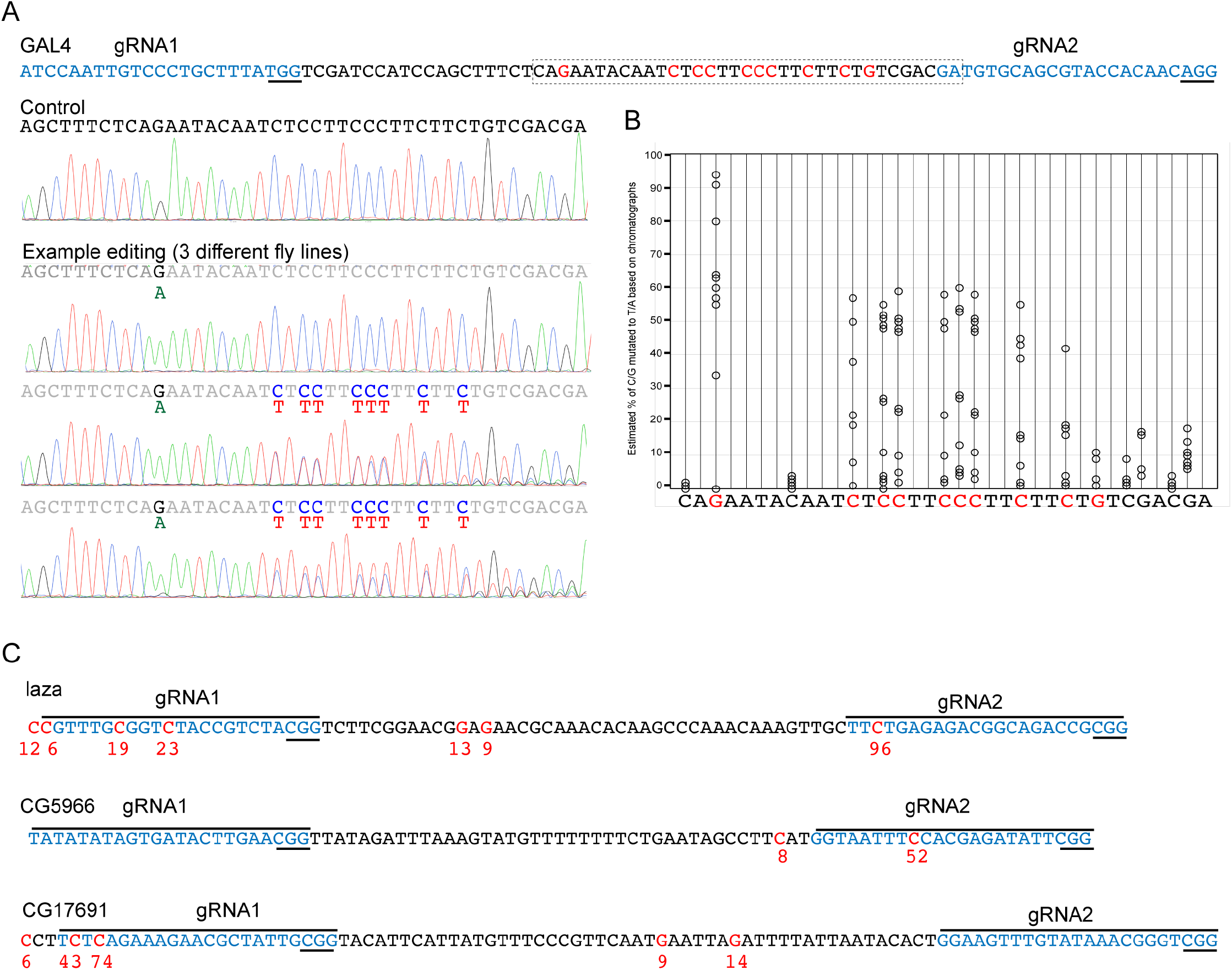
Expanded targeting observed when using closely spaced guide RNAs. **A)** The location of the two gRNAs in blue are shown on the GAL4 sequence. Residues changed by the DBE2 base editor are highlighted in red. Example wild-type and base-edited chromatographs from three different GAL4 fly lines are shown. Base editing occurred primarily outside the gRNA target. **B)** The EditR program was used to calculate the percentage of base-editing occurring for each base in the boxed region in **A. C)** Base-editing occurring between 3 additional genomic loci targeted by two closely spaced guide RNAs. The residues found to be mutated are in red. The number under the residue indicates the estimated percentage of conversion at that residue as calculated by EditR.

The data presented here indicate the accumulated editing that occurred throughout the hundreds of thousands of cells that make up an adult fly. It is likely that most base editing occurred early in development coinciding with expression of DBE2 from the *Actin5C* promoter. Once a mutation is induced in the gRNA target site (such as during early developmental), that site should no longer be a target for the gRNA, and no additional editing should occur. However, the high rates of editing seen here (multiple residues >50%) suggests base editing can occur at multiple cytosine residues during the same targeting event; this is consistent with *in vitro* oligomer assays of BE2 in mammalian cells (Komor et al., 2016).

In recent years a number of updates or additions to base editors have been released. To increase editing efficiency and reduce indels, BE3 was modified to variant BE4 which contains a longer linker between cytidine deaminase and nickase dCas9 and two copies of UGI (Komor et al., 2017). To broaden the editing window beyond the gRNA target sequence, the BE-PLUS system used dCas9 as a scaffold for 10 copies of the GCN4 peptide, which in turn could be complexed *in vivo* to a single chain variable fragment bound to base-editing molecular machinery (Jiang et al., 2018). In addition, alternative strategies for inducing mutations, such as the prime-editing technique (Bosch et al., 2021), are also effective in *Drosophila*. Base editing tools could be further paired with binary expression systems to limit CRISPR/Cas9 editing to specific adult tissues, such as the eye, for F1 mutagenesis screens. DBE2, and the utilization of other base editing reagents, further enrich the genomic engineering toolbox in *Drosophila*.

## Materials and Methods

### Cloning of pattB-Actin5C-DBE2 (pActin5c-CD-dCas9m4-UDI-attB, Addgene# 104879)

*pWallium-dCas9-VPR* (containing homo sapien codon optimized dCas9m4; Addgene# 78897 (Lin et al., 2015)) was digested with *NdeI*/*EcoRI*, and a 3.3 kB fragment isolated. This fragment was used to replace the *NdeI*/*EcoRI* fragment from *pCMV-BE2* (Addgene#73020 (Komor et al., 2016)) digested with *NdeI* and *EcoRI* using a 3-way ligation reaction (Rapid DNA ligation kit, Roche). This generated the intermediate plasmid *pCMV-BE2+dCas9m4*. The *BE2+Cas9m4* insert was PCR amplified and cloned using in-Fusion (Clontech) into the *pActin5c-attB-RFP* vector isolated by digestion of *pActin5c-Cas9* (Addgene# 62209 (Port et al., 2014)) with *EcoRI*/*KpnI*. The final plasmid was sequence verified.

### Fly Stocks

Wildtype flies were *IsoD1 (w*^*1118*^). The *GAL4* lines used were *ppk-GAL4* on III (BS#32079) and *NP2222-GAL4* on II (BS#112839). The two GAL4 gRNAs to target GAL4 were from GAL4>QF2 HACK Donor lines QF2^G4H^ 25C1 on II (as found in BS#66488) and QF2^G4H^ 65E3 on III (as found in BS#66495). The TRiP stocks, along with the gRNA target sequences, are summarized in the gRNA section below.

The GAL4 stocks to test *Actin5C-DBE2* were generated as follows. QF2^G4H^ 25C1 (DsRed+) /CyO; Dh/TM6B males were crossed to Pin/CyO; ppk-GAL4 (w+) virgins. Male and female progeny of genotype QF2^G4H^ 25C1/CyO; ppk-GAL4(w+)/TM6B were used to establish a stable stock. NP2222-GAL4/CyO; Dh/TM6B males were crossed to Pin/CyO; QF2^G4H^ 65E3/TM6B virgins, and progeny of genotype NP2222-GAL4/CyO; QF2^G4H^ 65E3/TM6B were used to establish a stable stock. To test the *Actin5C-DBE2*, the above stocks were crossed to *Actin5C-DBE2* on X virgin females, and F_1_ male progeny were analyzed for base editing.

### Generation of Actin5C-DBE2 transgenic flies (BS# TBD)

Transgenic insertion of *pattB-Actin5C-DBE2* onto the X chromosome was performed by PhiC31 integrase mediate transgenesis into docking strain *M{3xP3-RFP*.*attP}ZH-2A* located at cytological location 2A3 (Bischof et al., 2007). Injections were conducted by Rainbow Transgenic Flies Inc (Camarillo, California).

### Guide RNAs

A collection of transgenic gRNAs inserted at *attP40* (located at cytological location 25C6) were used from the Harvard Drosophila RNAi Screening Center (DRSC) and Transgenic RNAi Project (TRiP) collection (https://fgr.hms.harvard.edu/fly-in-vivo-crispr-cas). gRNAs were expressed from the *pCFD3* vector (single gRNA) or the *pCFD4* vector (two gRNAs) as listed below. Genomic targets were selected for those least likely to cause adverse effects if mutated, such as those in 5‘UTR sites targeted for gene activation using dCas9-VPR (Lin et al., 2015).

The fly stocks and gRNAs leading to base editing were (Bloomington Stock #, gRNA vector, gene target, gRNA sequence):

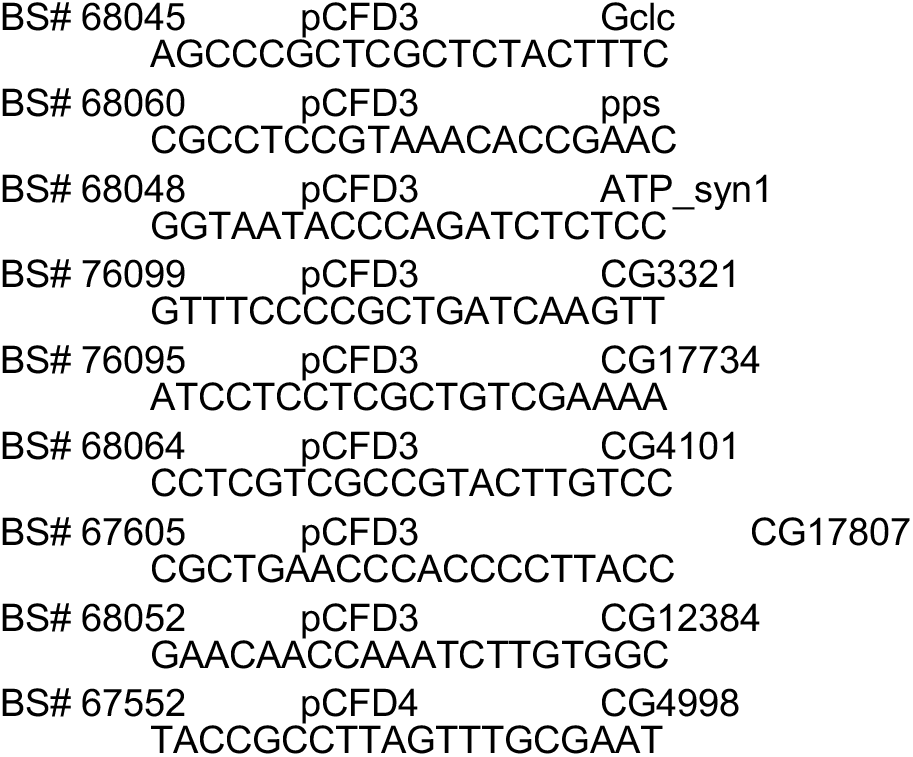

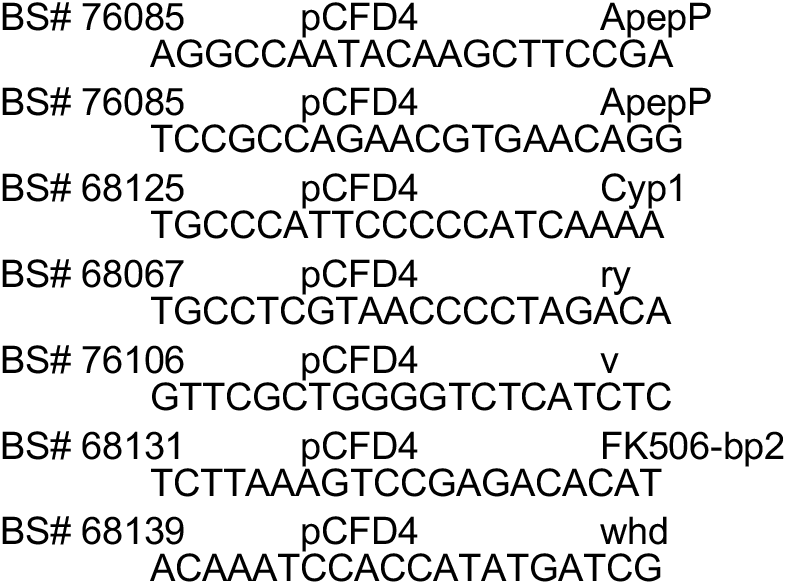

The fly stocks and gRNAs not leading to base editing were:

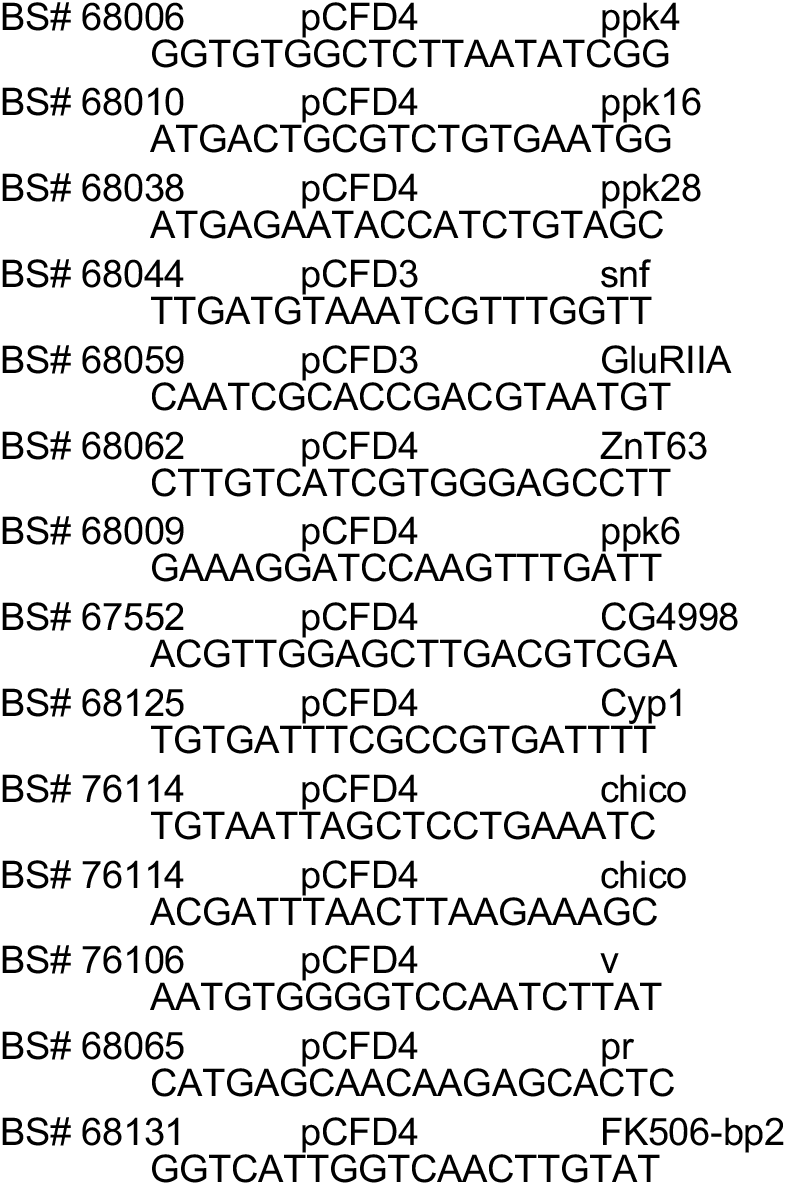

Note: the ability of these gRNAs to induce mutations using other Cas9 sources was not verified. It could be possible that mutation of these target sites might lead to cell death or reduced viability, and so might have been selected against during adult development.

The gRNA for GAL4 were from the GAL4>QF2 HACK donor line (Lin and Potter, 2016)

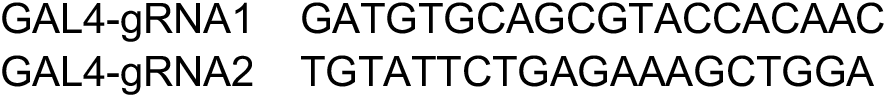

### Sequence analyses of gRNA targets

Genomic DNA surrounding the guide RNA targets was PCR amplified and sequenced. 10 F1 male flies were used as the source of genomic DNA as prepared by the QIAGEN DNeasy Blood & Tissue Kit (Qiagen Catalog # 69504). To control for genomic polymorphisms that might be misinterpreted as base editing or which mutated the target site, the target site was also PCR amplified and sequenced from control progeny containing a guide RNA targeting a different genomic location.

The following oligos were used for PCR amplification and sequencing:

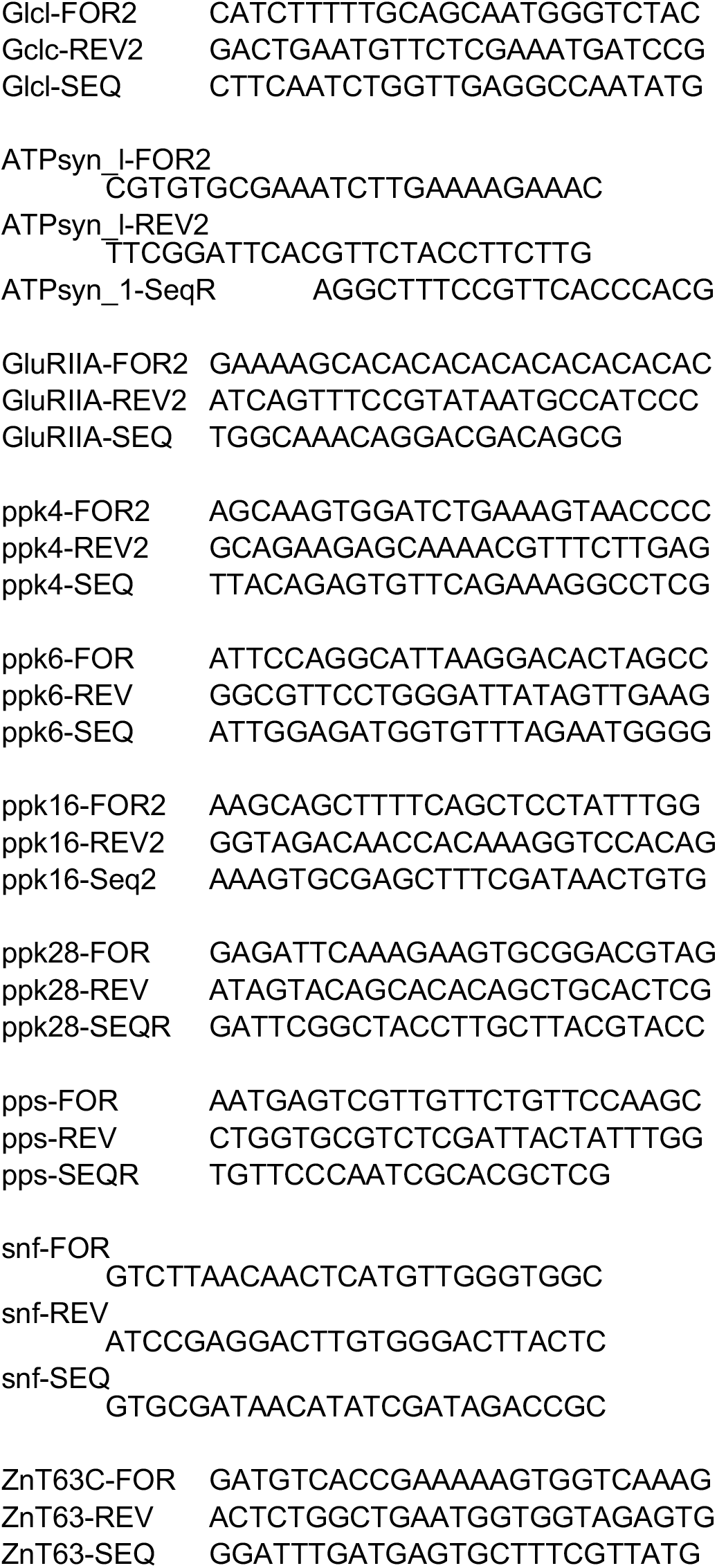

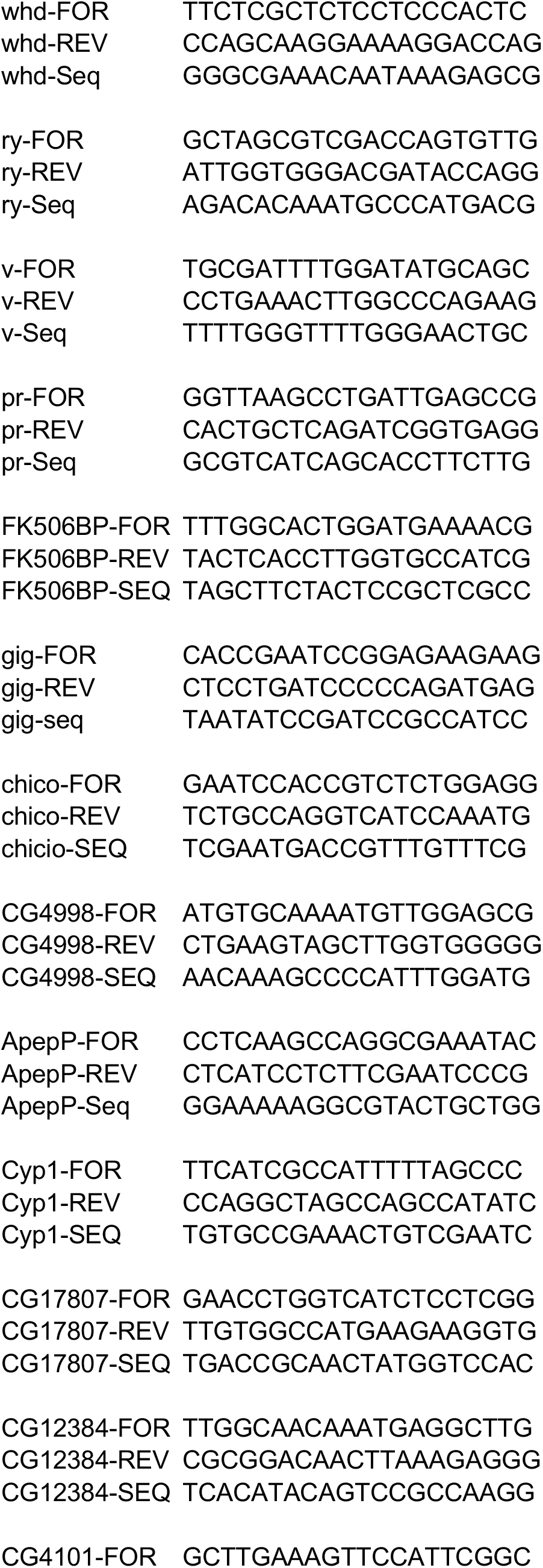

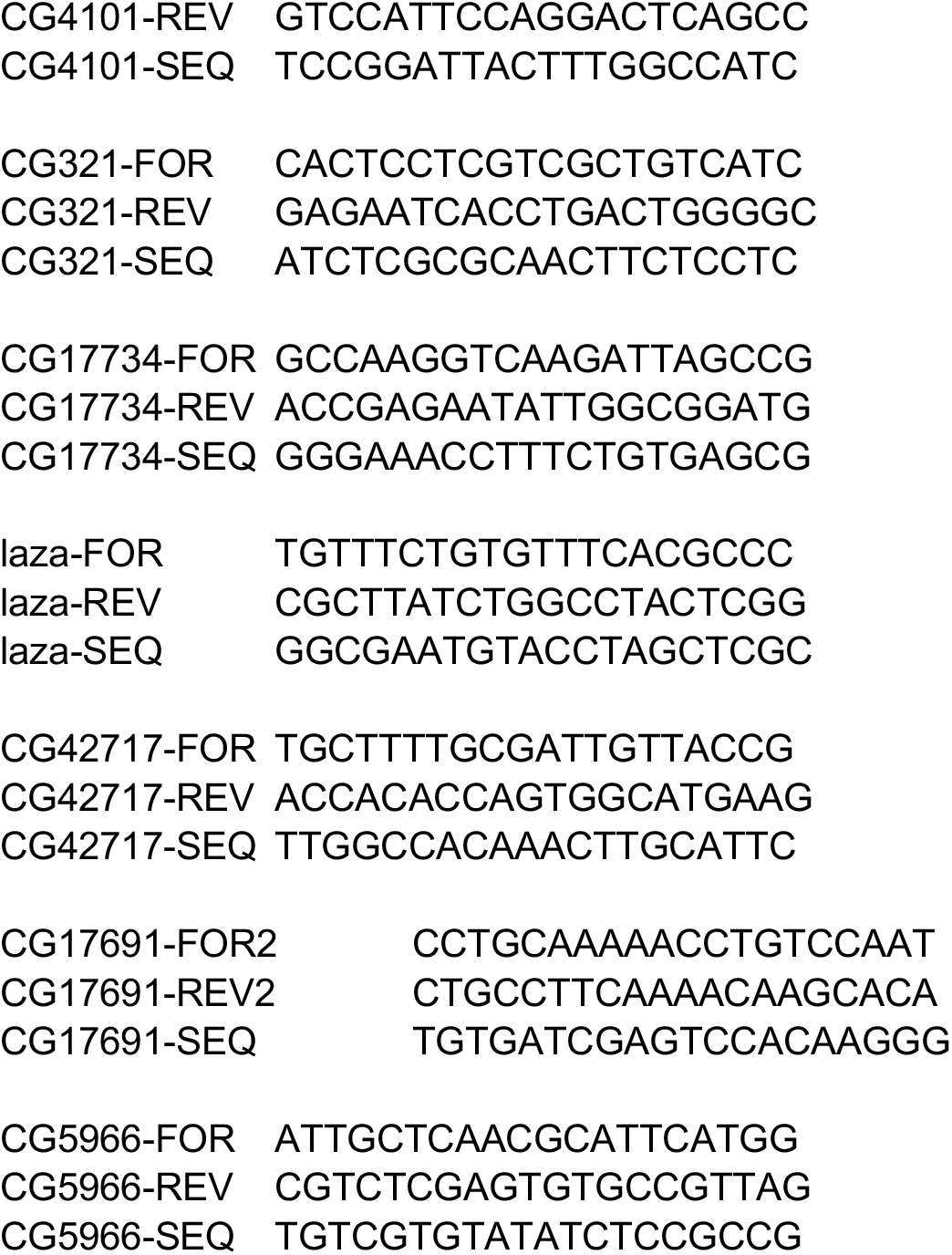

The following target sites were not included in the analyses due to polymorphisms in the target site that would disrupt at least one nucleotide in the gRNA target:

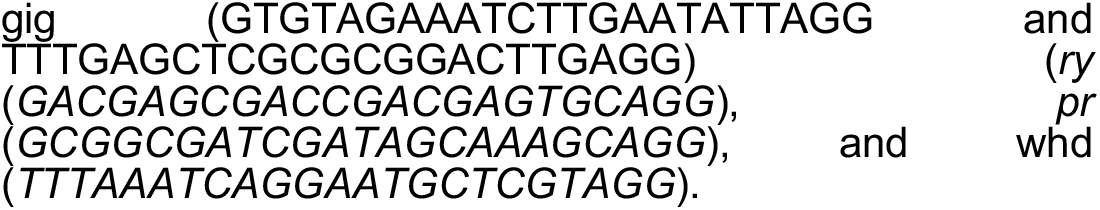

### Chromatogram analyses of Sanger sequencing files

The EditR program (http://baseeditr.com/)(Kluesner et al., 2018) was used to estimate the percentage of each base pair in the target sequences. The EditR program quantifies the area under each nucleotide peak at each position, and highlights regions likely targeted by base editing.

## Author Contributions (CRediT Taxonomy)

Conceptualization, C.J.P.; Methodology, C.J.P. and E.M.; Formal Analysis, C.J.P.; Investigation, E.M.; Writing – Original Draft, C.J.P.; Writing – Review & Editing, E.M. and C.J.P.; Visualization, C.J.P; Funding Acquisition, C.J.P.; Supervision, C.J.P.

The authors declare no conflict of interest.

## Acknowledgments

We thank Katie Robinson for assistance in conducting PCR analyses, and the Bloomington Drosophila Stock Center (NIH P40OD018537) for fly lines. This work was partly supported by the *NIH NIDCD (R01DC013070, CJP)*.

